# Frequency-domain identification of photosynthetic regulation under fluctuating light

**DOI:** 10.64898/2026.05.06.722921

**Authors:** Ladislav Nedbal

## Abstract

Plant photosynthesis operates under naturally fluctuating light, yet its dynamic responses across timescales remain incompletely understood. Here, we apply sinusoidal light modulation as a controlled periodic input and analyze the response in the frequency domain, enabling quantitative system identification of photosynthetic dynamics.

Using a minimal biochemical model of photosynthetic electron transport and regulation, we show that photosynthetic performance under fluctuating light differs systematically from that under constant illumination, even when the mean photon flux density is identical. Large-amplitude oscillations generate higher harmonics and alter time-averaged chlorophyll fluorescence, oxygen evolution, and non-photochemical quenching (NPQ), demonstrating that fluctuating light acts not merely as a perturbation but as a distinct physiological regime.

For sufficiently small perturbations, the system behaves approximately linearly and can be characterized by transfer functions and Bode plots. We identify two dynamic regimes separated by a characteristic timescale of approximately 10 s. In the high-frequency domain, the response is governed by constitutive photochemical processes and reflects local steady-state properties, including the redox state of the plastoquinone pool. In the low-frequency domain, adaptive regulatory feedback dominates, particularly NPQ, which reshapes both the amplitude and phase of the photosynthetic response.

Characteristic frequency-response features, including gain transitions and phase extrema, provide direct information about physiologically relevant quantities such as effective relaxation times and regulatory coupling strengths. We further introduce the concept of regulation fingerprints, defined as ratios of transfer functions between regulated and unregulated systems. These fingerprints reveal distinct spectral signatures of fast PsbS-dependent and slower zeaxanthin-dependent NPQ, enabling their quantitative separation and providing experimentally testable predictions for regulatory dynamics.

Together, these results establish frequency-domain analysis as a general framework for probing, identifying, and testing the dynamic regulation of photosynthesis under fluctuating light. More broadly, they suggest that fluctuating illumination, often regarded as experimental noise, can instead serve as a structured probe of photosynthetic function in both laboratory and field environments.

## 1. Introduction

### 1.1 Biological problem: photosynthesis under fluctuating light

Photosynthetic yield, productivity, and stress resilience depend not only on the mean intensity of photosynthetically active radiation but also on its temporal variability. In natural, agricultural, and biotechnological environments, light fluctuates across multiple timescales and a wide range of amplitudes—from rapid sunflecks and canopy movements to slower diel and environmental changes (Kaiser et al., 2018; Slattery et al., 2018; Table 1 in Niu et al., 2025). These fluctuations strongly influence carbon assimilation, energy dissipation, and photoprotective regulation (Kaiser et al., 2018).

Importantly, photosynthetic responses to fluctuating light are not generally equivalent to those under constant illumination with the same mean photon flux density. Nonlinear interactions between photochemistry and regulation can alter time-averaged photosynthetic performance, making fluctuating light a distinct physiological regime rather than a simple perturbation around steady-state behavior.

Most laboratory studies, however, rely on constant illumination or simple transitions between darkness and steady light. While such approaches provide high experimental control, they do not directly address how a fully light-acclimated photosynthetic system responds to dynamic perturbations around its operating state. As a result, the connection between mechanistic insights obtained under controlled conditions and photosynthetic performance under naturally fluctuating light remains incomplete (Long et al., 2022).

### 1.2 A systems perspective enables frequency-domain analysis of photosynthesis

To address this gap, photosynthesis can be probed by applying controlled, time-dependent perturbations and analyzing the resulting responses. In this work, sinusoidal light modulation is employed as a well-defined periodic input, enabling systematic characterization of photosynthetic dynamics in the frequency domain.

From this perspective, photosynthesis can be viewed as a nonlinear input–output system operating around a light-acclimated dynamic steady state (Alon, 2007; Klipp et al., 2016). Controlled oscillatory forcing, therefore, provides a means of probing the characteristic timescales, feedback strengths, and interactions that govern system behavior, analogous to system identification approaches widely used in engineering and systems biology.

Although the system comprises individual redox components, molecular complexes, and metabolite pools, its response to oscillatory light emerges from their collective interactions. Dynamic features observed under fluctuating light, including harmonic structure, nonlinear response amplitudes, and changes in dynamical stability (Section 2.1), cannot be attributed to any single component but arise from system-level coupling, conservation constraints, and feedback regulation.

Such distributed behavior is naturally addressed through mathematical modeling, which integrates physical, biochemical, and regulatory processes within a quantitative framework (Kitano, 2002). Mechanistic models, therefore, provide a means to identify the dynamical principles underlying photosynthetic regulation and its responses to fluctuating environments.

### 1.3 Strategy: combining mechanistic modeling with frequency-domain analysis

This study employs a minimal biochemical model of photosynthetic electron transport that captures regulatory nonlinearities across timescales ranging from milliseconds to thousands of seconds (Niu et al., 2025). The model deliberately excludes ultrafast photophysics and long-term acclimation, focusing instead on the intermediate dynamical regime in which feedback regulation, energy balance, and nonlinear responses dominate system behavior. Within this range, it preserves the essential feedback structure required to reproduce experimentally observed transient, oscillatory, and frequency-dependent responses (Niu et al., 2025).

An important advantage of this framework is that it enables systematic exploration of parameter space beyond experimentally accessible conditions. By varying rate constants, stoichiometries, and regulatory strengths, the model identifies the processes responsible for specific dynamical signatures, including harmonic structure, nonlinear gain, and changes in dynamical stability. Comparison between model predictions and experiments can then be used to test competing mechanistic hypotheses and guide further model refinement (Box, 1976).

Under constant illumination, the system relaxes to a stationary steady state. In contrast, periodic forcing drives it toward a periodic steady state, or limit cycle, determined jointly by the forcing frequency and the system’s intrinsic kinetic rates (Strogatz, 2015). The relationship between these timescales gives rise to distinct dynamic regimes. Small-amplitude oscillations enable linear transfer-function analysis and system identification.

At the same time, larger perturbations reveal nonlinear behavior through harmonic generation, changes in dynamical stability, and shifts in time-averaged photosynthetic performance.

Here, we combine sinusoidal light forcing, mechanistic modeling, and frequency-domain analysis to investigate how feedback regulation, energetic constraints, and multiple characteristic timescales shape photosynthetic dynamics under fluctuating light. We show that the system separates naturally into constitutive and regulatory dynamic domains (Nedbal and Lazár, 2021), that fluctuating illumination can modify time-averaged photosynthetic performance relative to constant light, and that individual regulatory mechanisms possess distinct spectral signatures that can be used for quantitative system identification. The following sections first analyze nonlinear responses to large-amplitude forcing and then use small-amplitude perturbations to resolve the constitutive and regulatory processes governing photosynthetic dynamics.

## 2. Results

### 2.1 Nonlinear Dynamics Under Large-Amplitude Forcing

#### 2.1.1 Time-domain signatures reveal nonlinear photosynthetic dynamics

Large-amplitude light modulation drives photosynthesis into a nonlinear response regime. Earlier studies have shown that chlorophyll fluorescence (ChlF) under sinusoidally modulated light contains a limited number of higher-order harmonics and, in higher plants, exhibits resonance-like behavior at periods of tens of seconds (Nedbal and Březina, 2002).

Similar phenomena have been observed in nonphotochemical quenching (NPQ), CO₂ assimilation, the Δ820 signal, and other photosynthetic observables across diverse organisms (Nedbal et al., 2003, 2005; Nedbal and Lazár, 2021; Lazár et al., 2022; Fuente et al., 2024). In *Arabidopsis thaliana*, the harmonic structure of ChlF responses has been shown to depend strongly on mutations affecting regulatory processes and cyclic electron transport (Niu et al., 2023, 2024, 2025).

In the present simulations, nonlinear behavior is evident directly in the time domain. State-space trajectories deviate from the elliptical loops expected for linear sinusoidal dynamics, indicating nonlinear interactions among the underlying processes. Correspondingly, the fluorescence waveform shown for the long-period oscillations in Fig. 1 (top right panel) and analyzed in detail in Fig. 2 departs from a purely sinusoidal shape. In the frequency domain, these distortions manifest themselves as higher-order harmonic components, providing a direct signature of nonlinear system behavior. The presence and relative strength of these harmonics therefore establish the boundary between genuinely nonlinear dynamics and the approximately linear regime analyzed in the following sections.

**Fig. 1.**
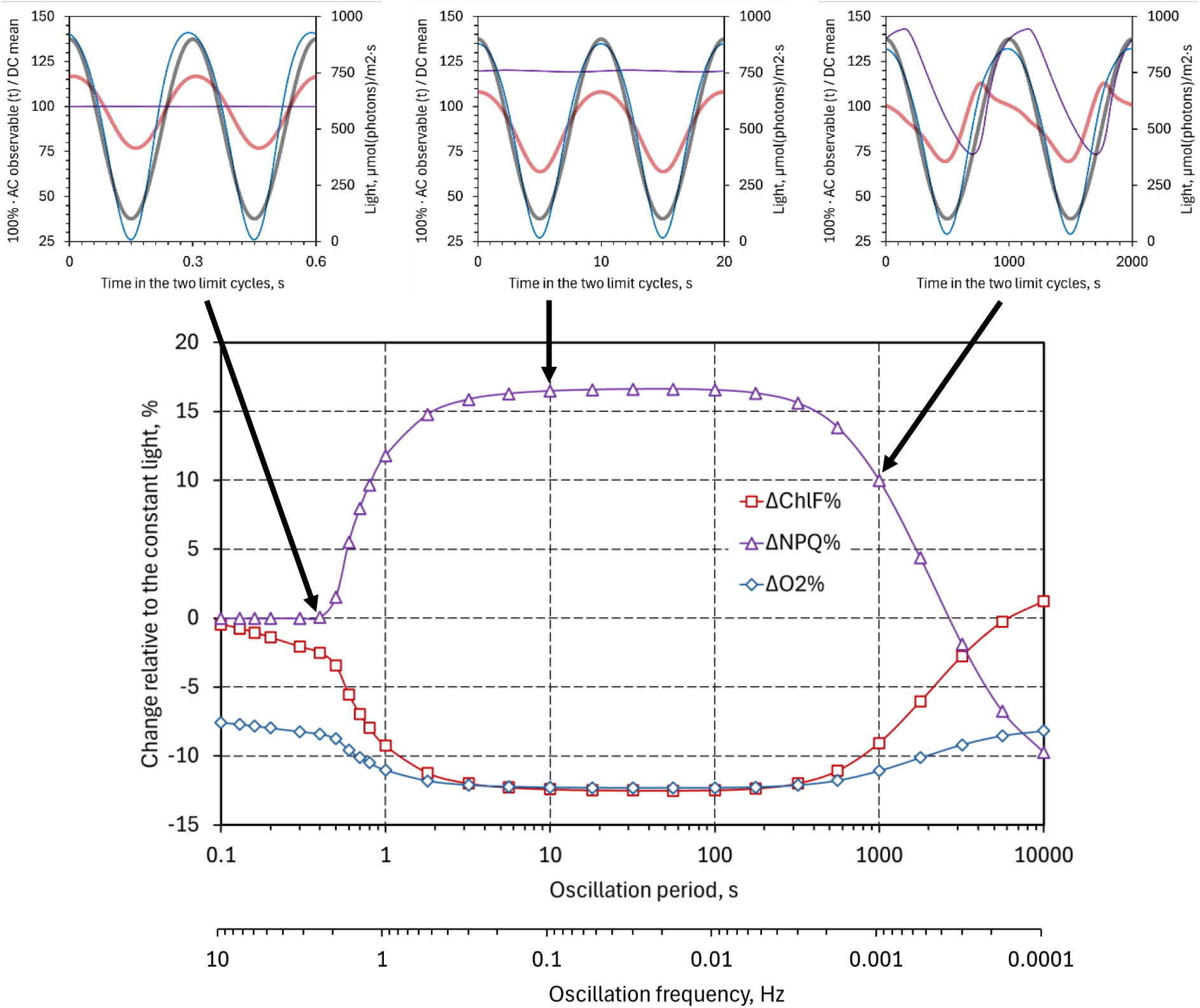
Effects of oscillating light on time-averaged photosynthetic observables. The top panels show relative changes in chlorophyll fluorescence yield (red), oxygen evolution rate (blue), and nonphotochemical quenching (NPQ, violet) under sinusoidal light oscillating between 100 and 900 µmol photons m⁻² s⁻¹ with periods of T = 0.3 s (left), 10 s (middle), and 1000 s (right). The responses are expressed as percentage changes relative to constant illumination with the same mean photon flux density. The bottom panel summarizes these changes over oscillation periods ranging from 100 ms to several hours. The effects of oscillating light are small at very short and very long periods but become pronounced at intermediate periods, where regulatory processes cannot fully equilibrate within a single oscillation cycle.

**Fig. 2.**
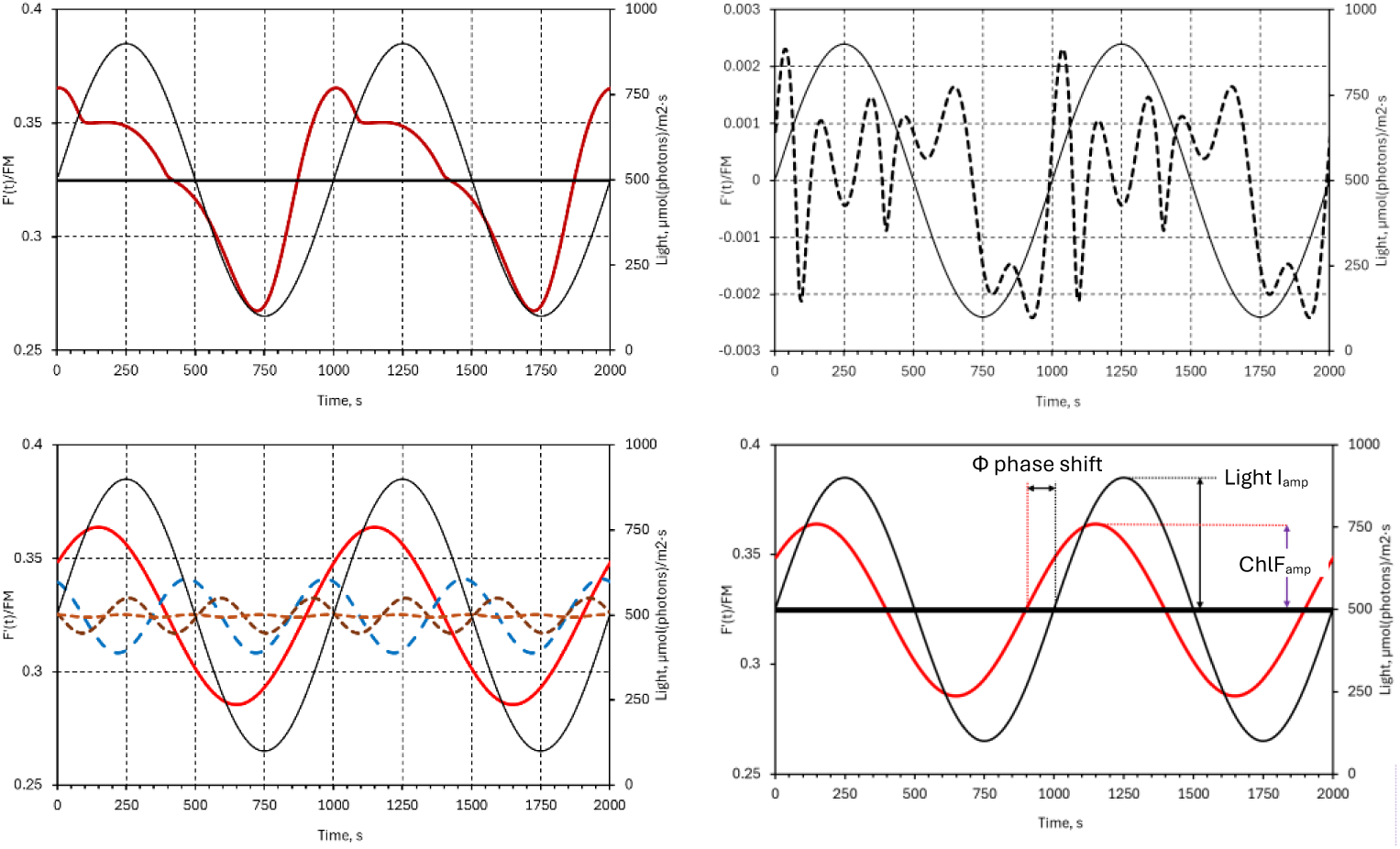
Harmonic decomposition of the simulated chlorophyll fluorescence (ChlF) response to sinusoidally oscillating light. The incident light (black line in all panels) oscillates between 100 and 900 µmol photons m⁻² s⁻¹ with a period of T = 1000 s (f = 1 mHz). The top-left panel shows the simulated ChlF response (dark red). The bottom-left panel presents its decomposition into the fundamental and the first three higher-order harmonic components. The top-right panel shows the residual difference between the full simulation and the four-harmonic reconstruction, demonstrating that the response is well represented by a compact harmonic spectrum. The bottom-right panel illustrates the amplitude and phase of the fundamental harmonic, which form the basis of the subsequent transfer-function analysis.

#### 2.1.2 Harmonic decomposition provides a compact representation of nonlinear responses

The nonlinear fluorescence signal can be decomposed into harmonic components using Fourier analysis (Fig. 2). Despite the visible waveform distortions, the response is well represented by a constant (mean) component, the fundamental harmonic at the forcing frequency, and only a limited number of higher-order harmonics. The residual remaining after subtraction of these components is negligible, indicating that the nonlinear dynamics are characterized by a compact harmonic spectrum rather than by broadband or irregular fluctuations. This suggests that the underlying regulatory processes operate through a small number of dominant dynamical modes.

The fundamental harmonic is of particular importance because it captures the component of the response oscillating at the forcing frequency and is fully characterized by its amplitude and phase relative to the input. These quantities provide the basis for the transfer-function analysis developed in the following sections. Higher-order harmonics, in contrast, quantify departures from linear behavior and reflect nonlinear interactions between light-driven excitation and regulatory processes. Their presence and relative amplitudes therefore provide quantitative signatures of the underlying nonlinear dynamics.

Harmonic decomposition thus provides a compact representation of photosynthetic responses under fluctuating light, linking waveform distortions observed in the time domain to the spectral structure revealed in the frequency domain. It also establishes a natural transition from the nonlinear regime considered here to the linear system-identification framework developed below, in which the fundamental harmonic becomes the primary descriptor of system dynamics.

#### 2.1.3 Fluctuating light modifies time-averaged photosynthetic performance

A further consequence of nonlinear dynamics is that photosynthetic responses under oscillating light differ from those under constant illumination, even when the mean photon flux density is identical. In nonlinear systems, the average response to a fluctuating input generally differs from the response to the average input. Time-averaged observables, such as oxygen evolution rate, chlorophyll fluorescence yield, and non-photochemical quenching (NPQ), therefore depend not only on the mean irradiance but also on its temporal structure.

Figure 1 illustrates this effect across a broad range of oscillation periods. For very rapid fluctuations, the photosynthetic apparatus effectively experiences the time-averaged irradiance, and the responses approach those observed under constant light. For very slow modulation, the system follows the instantaneous light level quasi-statically. Between these limits, regulatory processes cannot fully equilibrate within a single oscillation cycle.

In this intermediate regime, nonlinear interactions between light-driven excitation and regulatory feedback produce systematic shifts in time-averaged observables despite identical mean photon input. In the simulations shown in Fig. 1, photosynthetic performance—represented by oxygen evolution rate and chlorophyll fluorescence yield—is reduced under oscillating light relative to constant illumination over a broad range of oscillation periods, from approximately 1 s (1 Hz) to 1000 s (1 mHz). This interval coincides with the timescales of many natural light fluctuations (Kaiser et al., 2018; Slattery et al., 2018; Table 1 in Niu et al., 2025).

These results demonstrate that fluctuating light constitutes a distinct physiological regime rather than a simple perturbation around steady-state behavior. The observed deviations arise from the nonlinear dependence of photosynthetic output on input, whereby the response to average irradiance differs from the average response to fluctuating illumination. Frequency-domain analysis, therefore, provides a natural framework for quantifying how temporal variability itself shapes photosynthetic performance.

#### 2.1.4 Intermediate timescales exhibit the slowest convergence to periodic steady states

The period of oscillating light influences not only the steady-state response but also the time required for the photosynthetic system to adapt from constant illumination to a stable periodic state. The number of oscillation cycles required to reach this limit cycle, estimated from the chlorophyll fluorescence (ChlF) signal using a stroboscopic convergence criterion (Strogatz, 2015), is shown in Fig. 3. Because convergence depends on the ability of internal processes to adjust to the imposed forcing, it provides additional information about the characteristic timescales governing photosynthetic dynamics.

**Fig. 3.**
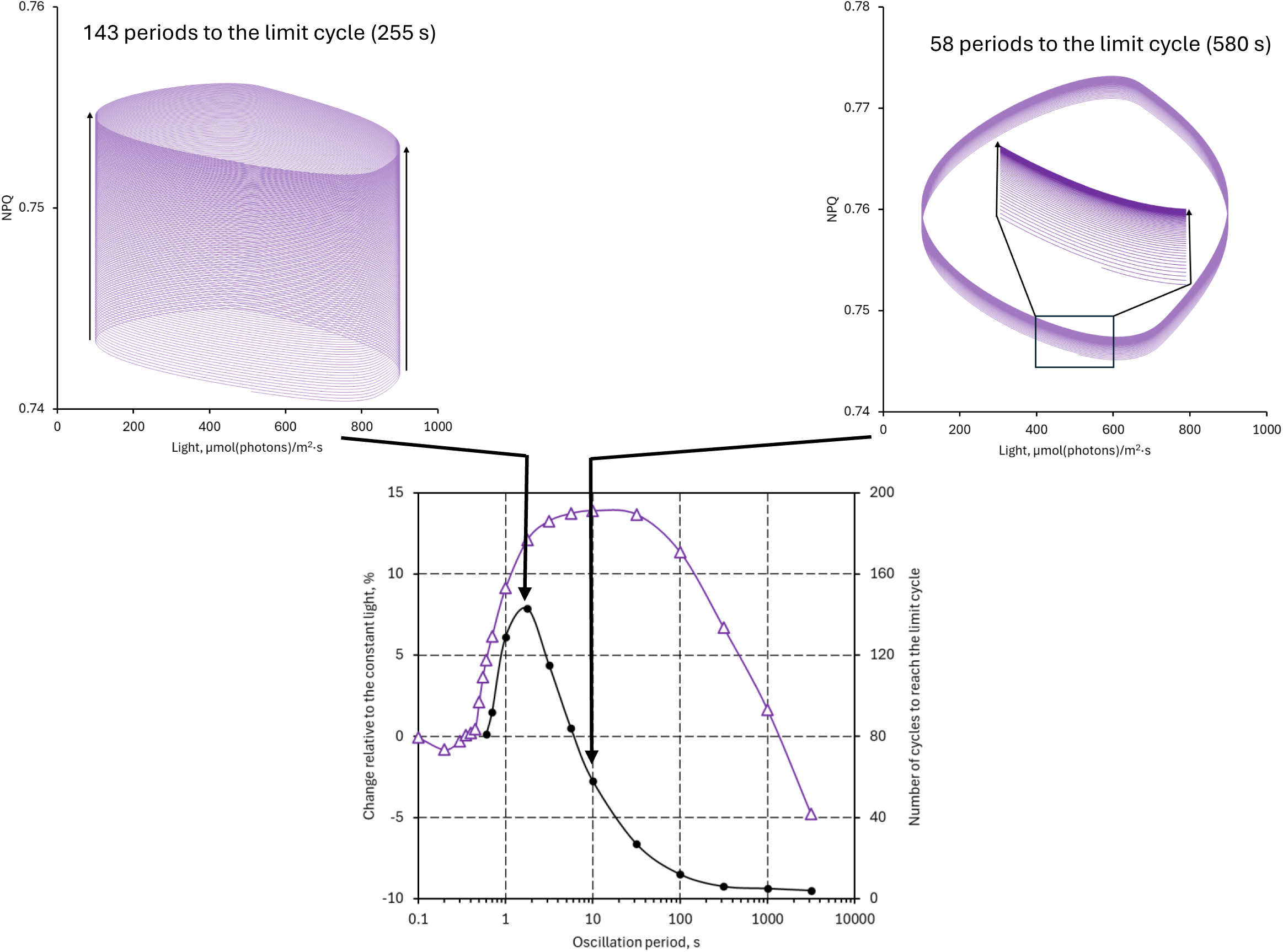
Convergence to periodic steady states under oscillating light. The bottom panel shows the number of oscillation cycles required for the photosynthetic system to converge from the steady state under constant illumination to a stable limit cycle under sinusoidal forcing of different periods. Convergence was quantified from the chlorophyll fluorescence (ChlF) signal using a stroboscopic criterion. The top panels illustrate representative convergence trajectories for forcing periods of T = 3.16 s (left, 143 cycles to convergence) and T = 10 s (right, 58 cycles to convergence). The slower convergence at intermediate periods reflects the mismatch between fast photochemical processes and slower regulatory responses, which cannot fully equilibrate within a single oscillation cycle. The maximum in convergence time occurs near the transition between constitutive and regulatory dynamics identified independently by the subsequent frequency-domain analysis.

A pronounced maximum in convergence time is observed at forcing periods of a few seconds, below the dominant regulatory timescales. In this regime, fast photochemical processes respond within individual oscillation cycles, whereas slower regulatory processes cannot fully adjust to the rapidly changing conditions. Consequently, many cycles are required before the system reaches a stable periodic response.

The phase-space trajectories for T = 3.16 s and T = 10 s illustrate this behavior. Convergence is substantially slower at T = 3.16 s, whereas at T = 10 s the system approaches the limit cycle more rapidly as regulatory processes begin to track the forcing.

The convergence maximum occurs near the transition between constitutive and regulatory dynamics identified independently by the frequency-domain analysis.

The convergence dynamics therefore provide an additional manifestation of the separation of timescales governing photosynthetic regulation. Convergence time is not merely a property of the numerical simulations but a dynamical observable that reflects the interplay between fast photochemical reactions and slower regulatory feedback processes. The agreement between convergence behavior and the independently derived frequency domains further supports the interpretation that photosynthetic dynamics are organized around distinct constitutive and regulatory timescales.

#### 2.1.5 Low-amplitude forcing defines the linear identification regime

The degree of nonlinearity in the photosynthetic response can be quantified using total harmonic distortion (THD), which measures the combined strength of higher-order harmonics relative to the fundamental oscillation. Because higher harmonics arise from nonlinear interactions, THD provides a convenient measure of the departure from linear behavior. It depends on the background light intensity, the modulation amplitude, and the forcing period (Fig. 4).

**Fig. 4.**
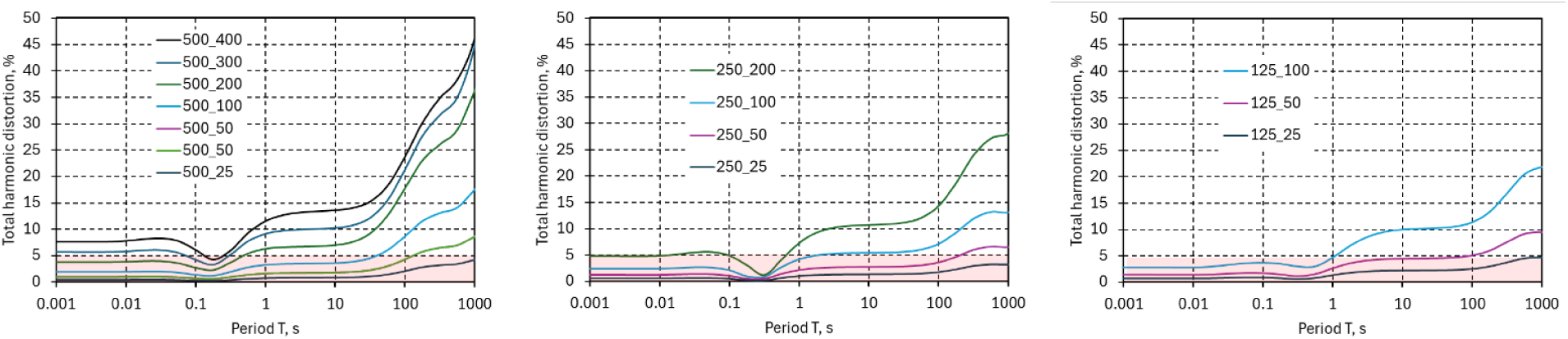
**Total harmonic distortion (THD)** of the chlorophyll fluorescence (ChlF) signal as a function of oscillation period for background light intensities of 500 (left), 250 (middle), and 125 µmol photons m⁻² s⁻¹ (right). The numbers in the legends indicate the amplitudes of the light modulation. THD quantifies the contribution of higher-order harmonics relative to the fundamental oscillation and therefore provides a measure of nonlinear behavior. The shaded region at the bottom of each panel marks THD values below 5%, which are used here as an operational criterion for approximate linearity and for the applicability of linear transfer-function analysis.

The simulations show that nonlinearity decreases as modulation amplitude decreases. For sufficiently small perturbations, THD remains below approximately 5% over a broad range of oscillation periods. Under these conditions, the fluorescence response is dominated by the fundamental harmonics, while higher-order harmonics contribute only weakly. The system can therefore be approximated as linear and time-invariant around the operating point defined by the constant background illumination. The 5% threshold is used here as a practical criterion for identifying the regime in which linear system identification is applicable.

This transition from harmonic-rich nonlinear dynamics to responses dominated by the fundamental oscillation establishes the connection between nonlinear and linear descriptions of photosynthesis. Once higher-order harmonics become negligible, the fundamental harmonic provides a compact characterization of system dynamics through its amplitude and phase, thereby enabling the transfer-function and Bode-plot analyses developed in the following section.

### 2.2 Linear transfer-function identification

#### 2.2.1 Small perturbations enable frequency-domain system identification

Low-amplitude light oscillations produce approximately linear responses that are dominated by the fundamental harmonic (Fig. 4). Under these conditions, the response at each forcing frequency is fully characterized by its amplitude and phase relative to the input. The photosynthetic system can therefore be treated as a linear time-invariant system around its light-acclimated operating point and described by transfer functions that relate oscillations in incident light to those in the measured observables.

This approximation enables quantitative system identification in the frequency domain. Rather than probing transitions between distinct physiological states, the analysis characterizes the local dynamical properties of the system around its actual operating point, including the timescales and coupling strengths associated with photochemical and regulatory processes.

The resulting Bode plots reveal distinct dynamical domains associated with constitutive photochemical processes and slower regulatory mechanisms. As shown below, the observed gain and phase changes can be interpreted mechanistically in terms of plastoquinone (PQ) dynamics and the NPQ regulation (see Methods, Sections 5.5 and 5.6).

#### 2.2.2 Bode plots separate constitutive and regulatory photosynthetic dynamics

Primary photochemical processes and regulatory mechanisms jointly shape the response of chlorophyll fluorescence (ChlF) to sinusoidally modulated light under constant background illumination (Figs. 5 and 6). The gain characterizes the magnitude of the fluorescence response relative to the light modulation, whereas the phase describes its delay with respect to the forcing.

**Fig. 5.**
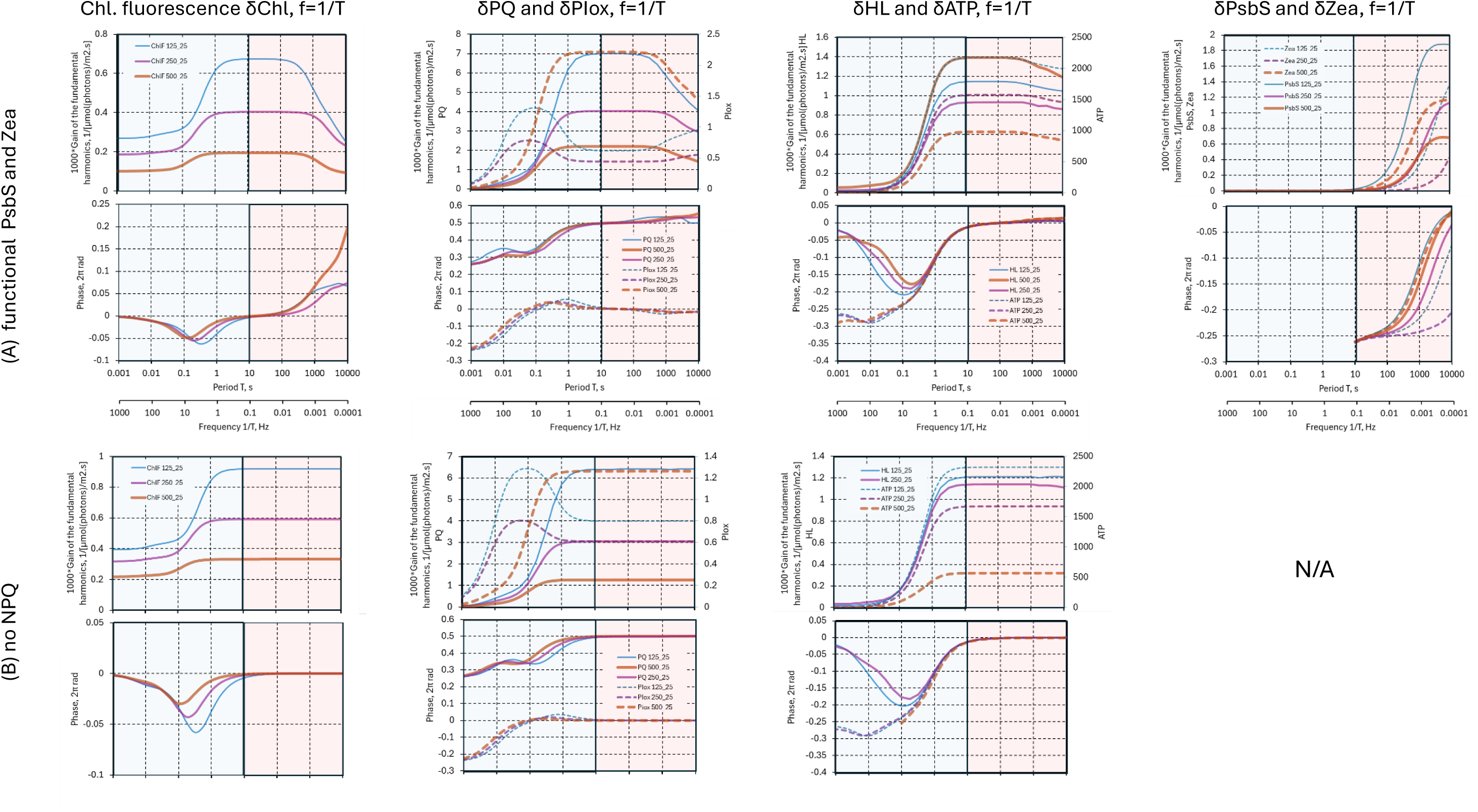
**Bode plots** of chlorophyll fluorescence (δChl), plastoquinone redox dynamics (δPQ), oxidized PSI donors (δPIox), lumen proton concentration (δHL), ATP (δATP), and the PsbS and Zea quenchers (δPsbS and δZea) simulated with the BDM2 model. Panel A shows simulations with both fast (PsbS-dependent) and slow (Zea-dependent) nonphotochemical quenching (NPQ) mechanisms active, whereas panel B shows simulations without NPQ regulation. The remaining panels illustrate how the underlying photochemical and regulatory variables contribute to the observed fluorescence dynamics. The simulations used the original parameterization of Niu et al. (2025). Colors indicate background light intensities of 125 (blue), 250 (magenta), and 500 (brown) µmol photons m⁻² s⁻¹. The modulation amplitude was fixed at 25 µmol photons m⁻² s⁻¹, maintaining total harmonic distortion below 5% and ensuring approximately linear system behavior. The shaded regions indicate the high-frequency constitutive (blue) and low-frequency regulatory (red) domains discussed in the text, demonstrating how different frequency ranges selectively probe photochemical and regulatory processes.

**Fig. 6.**
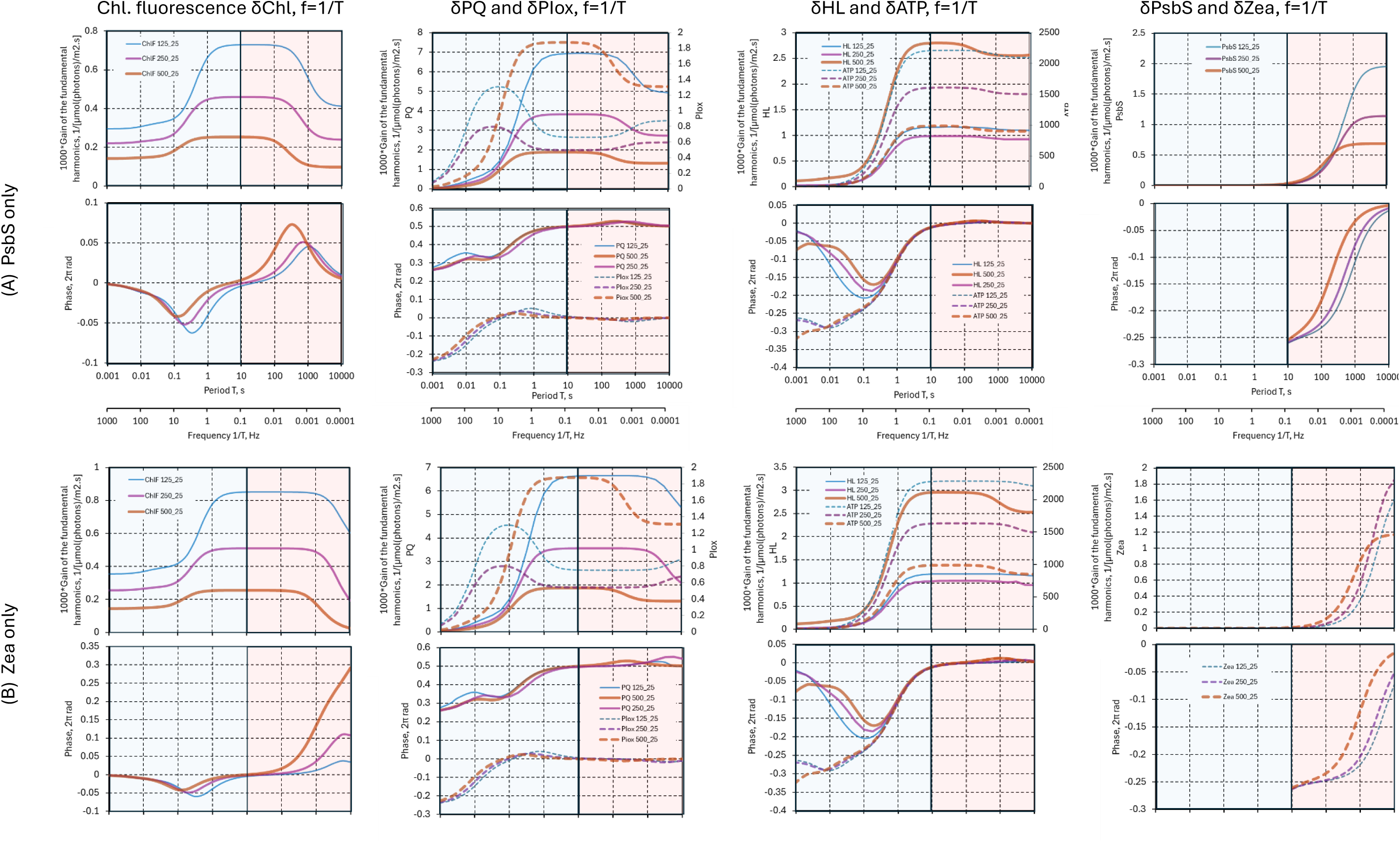
Bode plots illustrating the distinct spectral signatures of the fast PsbS-dependent and slow zeaxanthin (Zea)-dependent components of nonphotochemical quenching (NPQ) simulated with the BDM2 model. The panels show the responses of chlorophyll fluorescence (δChl), plastoquinone redox dynamics (δPQ), oxidized PSI donors (δPIox), lumen proton concentration (δHL), ATP (δATP), and the respective quenching variables. The simulations used the original parameterization of Niu et al. (2025). Colors indicate background light intensities of 125, 250, and 500 µmol photons m⁻² s⁻¹. The comparison reveals that the two regulatory mechanisms operate on distinct characteristic timescales and contribute differently to the frequency-dependent modulation of photosynthetic dynamics.

Figure 5A shows simulations performed using the parameter set of Niu et al. (2025), including the original kinetic description of the PsbS- and Zea-dependent NPQ components. The corresponding Bode plots reveal distinct frequency domains associated with constitutive photochemical processes and slower regulatory feedback. At high frequencies, the response is dominated by the intrinsic dynamics of electron transport, whereas at lower frequencies NPQ regulation increasingly shapes both the gain and phase characteristics.

The remaining panels display the frequency responses of the underlying model variables, including plastoquinone redox dynamics (δPQ), oxidized PSI donors (δPIox), lumen proton concentration (δHL), ATP dynamics (δATP), and the PsbS- and Zea-dependent quenchers.

Together, these internal variables provide a mechanistic basis for interpreting the fluorescence response and for identifying how photochemical and regulatory processes contribute to distinct dynamical regimes.

#### 2.2.3 Frequency-domain analysis links fluorescence signals to underlying mechanisms

The linear decomposition of the fluorescence signal (Methods, Section 5.3) provides a mechanistic interpretation of the frequency-dependent structure of the Bode plots in Figs. 5 and 6. In this framework, the fluorescence response is represented as the sum of a direct light-driven contribution, a plastoquinone-mediated constitutive component, and slower regulatory terms associated with PsbS- and zeaxanthin-dependent NPQ.

At all frequencies, the direct light-driven contribution affects the gain but does not introduce a phase delay. At intermediate periods (∼0.1–1 s), the PQ-mediated constitutive component becomes dominant. The delayed response of the plastoquinone pool produces the characteristic phase minimum observed in the Bode plots.

At longer periods (>10 s), the regulatory contributions associated with PsbS and zeaxanthin become increasingly important. These slower processes suppress the gain and introduce additional phase delays, reflecting the characteristic timescales of NPQ regulation.

The frequency-domain separation of these contributions therefore establishes a direct connection between observable fluorescence dynamics and the underlying biochemical processes governing photosynthetic regulation. Distinct features of the Bode plots, including gain plateaus, phase minima, and phase maxima, can thus be interpreted in terms of specific constitutive and regulatory mechanisms.

#### 2.2.4 High-frequency dynamics reflect constitutive photochemical processes

In the high-frequency domain (periods shorter than ∼10 s; blue-shaded region in Fig. 5), chlorophyll fluorescence dynamics are determined primarily by the incident light and the redox state of the plastoquinone pool (PQ). Other state variables influence the response only indirectly through their effects on PQ dynamics or on slowly varying parameters entering the fluorescence model.

The constitutive nature of this domain is demonstrated independently by simulations in which NPQ regulation is disabled (Fig. 5B). In the absence of regulatory feedback, the system still exhibits two gain plateaus separated by a single-phase minimum, characteristic of an effective lead–lag response that emerges from the interaction of processes operating on different timescales. Specifically, the response reflects the coupled dynamics of the PQ pool and the oxidized PSI donor side (PIox).

At periods shorter than ∼10 ms, variations in PQ become strongly suppressed and the fluorescence response is dominated by the direct light-driven contribution. The phase shift approaches zero and the gain converges toward a constant high-frequency limit determined by the local steady-state operating point (Methods, Section 5.5).

Thus, unlike the low-frequency domain analyzed below, which is governed by adaptive negative-feedback regulation, the high-frequency domain reflects constitutive charge-transfer dynamics within the photosynthetic electron transport chain. The apparent symmetry between short- and long-period fingerprints is therefore only phenomenological: the former arise from intrinsic redistribution dynamics, whereas the latter emerge from genuine regulatory feedback.

#### 2.2.5 Low-frequency dynamics are governed by NPQ regulation

The low-frequency domain (periods longer than ∼10 s; red-shaded region in Fig. 5) is dominated by regulatory processes, primarily the synergistic action of the PsbS- and zeaxanthin (Zea)-dependent components of non-photochemical quenching (NPQ). Their combined effect is illustrated in Fig. 5A.

The distinct spectral signatures and timescales of these regulatory mechanisms are shown in Fig. 6, where the contributions of PsbS and Zea can be analyzed separately. In this regime, the quenchers suppress oscillation amplitudes across multiple variables, including chlorophyll fluorescence (ChlF; left column), the plastoquinone redox state (PQ; middle-left column), and the lumen proton concentration and ATP (middle-right column).

The PSI donor side (PIox), however, exhibits a more complex dependence on background light intensity. At lower light levels (125 and 250 µmol photons m⁻² s⁻¹), activation of the quenchers reduces the oscillation amplitude at low frequencies. In contrast, at the highest light level (500 µmol photons m⁻² s⁻¹), the trend is reversed, and the amplitude of Piox oscillations increases as the quenchers become dynamically active.

These results show that NPQ regulation does not merely suppress fluctuations but actively reorganizes the dynamical relationships among photosynthetic variables. The frequency-dependent activation of NPQ therefore modulates both the amplitude and phase of photosynthetic responses under fluctuating light.

In contrast to the high-frequency domain, which reflects constitutive charge-transfer processes, the low-frequency domain provides direct access to the dynamics and interactions of regulatory feedback mechanisms.

#### 2.2.6 Regulation fingerprints distinguish fast and slow NPQ mechanisms

Regulatory processes shape photosynthetic dynamics across the frequency range, most prominently in the low-frequency domain (Figs. 5 and 6), while also indirectly influencing higher-frequency behavior through modulation of constitutive processes.

To isolate the dynamic contribution of individual regulatory mechanisms, we introduce regulation fingerprints as the logarithm of the ratio between transfer functions of regulated and unregulated systems:

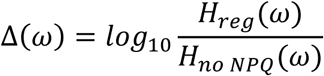

Specifically, we compare systems with PsbS- or zeaxanthin (Zea)-dependent regulation to the system without NPQ regulation (Fig. 7), mirroring experimental strategies in which mutants lacking individual NPQ components are used to isolate their dynamic contributions under fluctuating light (Niu et al., 2023).

**Fig. 7.**
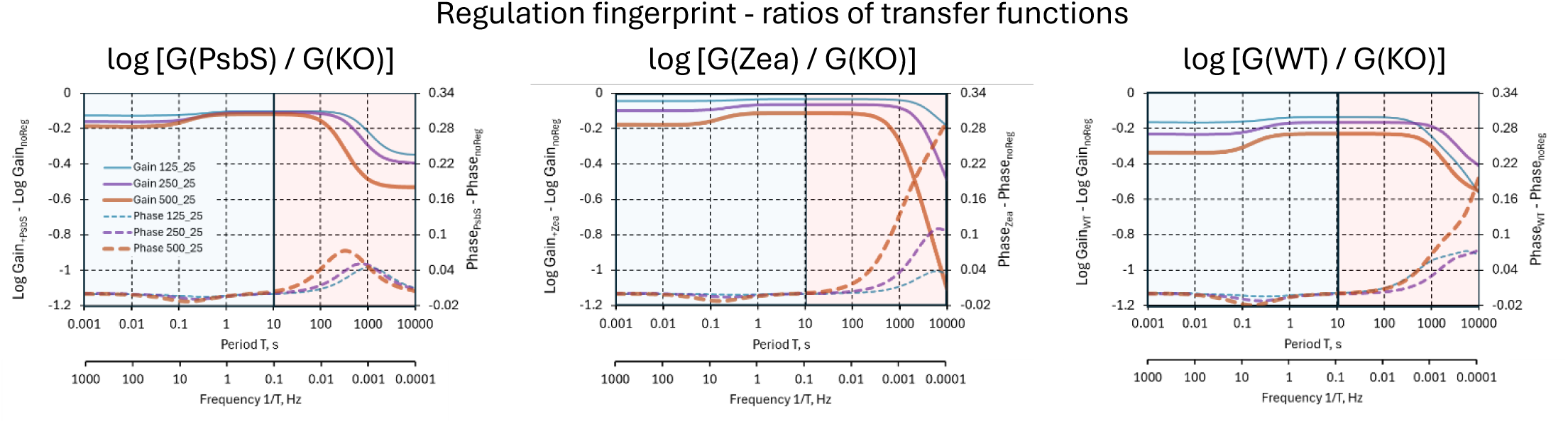
Regulation fingerprints. The transfer functions of BDM system with the PsbS regulation (left panel), with Zea regulation (middle panel) and with both regulations (right panel) are compared to the system without any regulation by plotting the logarithm of gain differences and differences of phase shifts.

### PsbS regulation

The PsbS fingerprint (Fig. 7, left panel) exhibits a transition in gain from a higher plateau at intermediate periods (tens to hundreds of seconds) to a lower plateau at longer periods approaching 10^4^ s. The phase difference is close to zero at both short and long periods but displays a pronounced maximum in the intermediate range.

The position of this phase maximum depends on light intensity, shifting from ∼1000 s at low light (125 µmol photons m⁻² s⁻¹) toward shorter periods (∼300–500 s) at higher light levels. These maxima occur near the inflection points of the gain transition and reflect a delayed negative-feedback response: at short periods, the regulation cannot respond, at intermediate periods, it introduces a phase lag, and at long periods, the system approaches quasi-steady adaptation.

From a control-theoretic perspective, this behavior is consistent with a feedback element with a characteristic timescale of several hundred seconds that decreases with increasing irradiance.

### Zea regulation

The Zea fingerprint (Fig. 7, middle panel) exhibits qualitatively similar behavior but is shifted toward longer periods, indicating slower dynamics. The characteristic features extend beyond the simulated range (>10^4^ s), consistent with the slower kinetics of zeaxanthin accumulation and relaxation.

The magnitude of both gain suppression and phase variation is larger than for PsbS, reflecting the stronger influence of this slower regulatory component. This suggests that Zea acts, in the model, as a deeper, longer-term modulation of energy dissipation compared to the faster PsbS-mediated response.

### Combined regulation (WT)

The wild-type (WT) fingerprint (Fig. 7, right panel) reflects a superposition of the PsbS and Zea contributions. The resulting response combines features of both fast and slow regulation, producing a multi-timescale feedback structure. The coexistence of these regulatory components generates a dynamical architecture that cannot be captured by a single characteristic relaxation time, emphasizing the inherently multi-timescale nature of photosynthetic regulation.

#### 2.2.7 Bode-plot features reveal physiologically relevant NPQ parameters

Beyond distinguishing fast and slow regulatory mechanisms, the dynamic fingerprints establish quantitative links between experimentally observable Bode-plot features and physiologically meaningful parameters of NPQ regulation. In particular, the PsbS fingerprint (Fig. 7, left panel) relates the position of the phase maximum and the transition region of the gain to the characteristic relaxation time of PsbS activation (Methods, Section 5.7).

The simulations show that increasing background illumination shifts the phase maximum toward shorter periods (Fig. 7). This behavior is consistent with enhanced lumen acidification, which increases the activation function *A*(〈*H_L_*〉) and thereby reduces the effective relaxation time of the PsbS regulatory pathway (Methods, Section 5.6). The characteristic timescale of NPQ regulation, therefore, depends not only on intrinsic kinetic parameters but also on the physiological operating point established by the prevailing irradiance.

The magnitude of the gain transition further reflects the effective coupling strength between lumen acidification and NPQ activation. Together, these spectral features provide experimentally accessible estimates of both relaxation times and regulatory gains without requiring direct measurements of internal state variables.

The same principles extend to the slower zeaxanthin-dependent pathway, whose characteristic frequencies occur at longer periods. In experimental systems, analogous information can be obtained by comparing wild-type plants with mutants lacking individual regulatory components, such as *npq4* and *npq1*, thereby linking frequency-domain fingerprints to specific molecular mechanisms.

These results demonstrate that frequency-domain measurements under fluctuating light provide a non-invasive approach for quantifying the dynamics of NPQ regulation and their dependence on environmental conditions. More generally, they establish Bode-plot features as experimentally measurable proxies for the physiological parameters governing photosynthetic feedback regulation.

## 3. Discussion

Photosynthesis operates under fluctuating light, yet its regulation is most often studied under steady-state or step-change conditions. The frequency-domain framework developed here provides a complementary perspective by probing the system around its operating state and resolving its dynamics across timescales.

A central outcome of this work is the separation of photosynthetic dynamics into frequency domains associated with different functional processes. Rather than reflecting a single mechanism, the observed responses arise from the interaction of fast photochemical reactions and slower regulatory feedback.

The frequency-domain representation makes this separation explicit and links observable signals to underlying processes.

Importantly, the analysis shows that frequency-response features are not merely descriptive but carry quantitative information. High-frequency behavior reflects local steady-state properties, while low-frequency dynamics encode regulatory timescales and coupling strengths. In this sense, oscillatory measurements provide access to system parameters that are difficult to obtain from steady-state experiments. The phase shifts observed in chlorophyll fluorescence directly reflect the delayed dynamics of regulatory activation. Increasing light intensity shifts the phase maxima toward shorter periods, consistent with enhanced lumen acidification and faster effective PsbS dynamics. Similar considerations are expected to apply to the slower Zea-dependent pathway, emphasizing that regulatory timescales depend not only on intrinsic kinetic parameters but also on the physiological operating state established by the incident light.

The concept of regulation fingerprints extends this idea by isolating the spectral contribution of individual regulatory components. The distinct signatures of PsbS- and zeaxanthin-dependent NPQ show that regulatory processes can be separated based on their characteristic timescales. Regulation fingerprints encode both the strength and kinetics of feedback, allowing different components of NPQ to be distinguished quantitatively. This provides a principled framework for using frequency-domain measurements to resolve the regulatory structure of photosynthesis and to relate complex fluorescence dynamics to specific biochemical mechanisms.

The concept of regulation fingerprints also provides a natural bridge between frequency-domain system identification and plant genetics. In experimental studies, mutants lacking individual regulatory components, such as the PsbS-deficient npq4 mutant, the violaxanthin de-epoxidase mutant npq1, or mutants affecting cyclic electron transport, have been used to isolate specific dynamical contributions under fluctuating light (Niu et al., 2023, 2024, 2025). The present framework suggests that such perturbations can be interpreted quantitatively in terms of changes in dynamic spectral signatures rather than solely through differences in steady-state or transient responses. Frequency-domain measurements may therefore provide a principled approach for relating genetic modifications to the characteristic timescales, coupling strengths, and feedback structures that govern photosynthetic regulation.

Frequency-domain measurements may also provide a powerful framework for model parameter identification. In the high-frequency constitutive domain, features such as the asymptotic gain, the position of the phase minimum, and their dependence on background illumination reflect local steady-state properties and characteristic electron-transfer timescales. At lower frequencies, the suppression of gain and the emergence of additional phase delays encode the activation dynamics of PsbS- and Zea-dependent regulation. In principle, characteristic quantities such as the effective relaxation time of PsbS or the coupling strength between lumen acidification and NPQ activation could therefore be inferred directly from Bode-plot features. Unlike conventional dark–light–dark protocols, which probe transitions between acclimation states, oscillatory measurements characterize the regulatory system around its physiological operating point and may provide complementary constraints for quantitative parameter estimation.

Importantly, these distinct spectral signatures provide an opportunity to test the model experimentally and, if necessary, to falsify it. Agreement between predicted and measured fingerprints would support the underlying mechanistic assumptions, whereas systematic deviations could indicate missing pathways, incorrect kinetic descriptions, or additional layers of regulation. In this sense, regulatory processes in photosynthesis can be characterized by their dynamic spectral signatures, analogous to the use of system identification methods in engineering.

At the same time, the present framework has limitations. The model is intentionally minimal and does not include all regulatory pathways or metabolic interactions. Additional processes, spatial heterogeneity, or coupling to carbon metabolism may introduce further dynamical features that are not captured here.

Likewise, the lead–lag behavior observed in the high-frequency constitutive domain should not be regarded as a universal property of photosynthetic electron transport but rather as an emergent consequence of the present parameterization. Different kinetic balances between electron donors and acceptors may produce qualitatively different frequency responses, including predominantly low-pass behavior. Extending the approach to more detailed models and experimental systems will therefore be necessary to assess the generality of the proposed dynamic fingerprints.

## 4. Conclusions and outlook

This study demonstrates that photosynthetic regulation can be quantitatively probed using frequency-domain analysis of responses to oscillating light. By combining controlled sinusoidal forcing with harmonic and transfer-function analysis, it becomes possible to extract physiologically relevant information, including characteristic regulatory timescales and coupling strengths, from chlorophyll fluorescence measurements. The resulting frequency fingerprints link observable dynamics to specific photochemical and regulatory processes and provide a framework for disentangling the contributions of multiple feedback mechanisms.

The results further show that fluctuating light constitutes a distinct physiological regime rather than merely a perturbation around steady-state behavior. Even under identical mean photon flux densities, nonlinear interactions between photochemistry and regulation modify time-averaged photosynthetic performance. Frequency-domain analysis, therefore, provides not only a description of dynamic responses but also a quantitative framework for understanding how temporal variability itself influences photosynthetic function.

A key practical implication is that small-amplitude light modulation offers a non-invasive approach to system identification in photosynthesis. In principle, this methodology can be implemented under laboratory, greenhouse, and field conditions, enabling characterization of photosynthetic regulation *in vivo* under realistic environmental fluctuations. Unlike conventional protocols based on dark acclimation and saturating flashes, frequency-domain measurements probe the system around its physiological operating point and benefit from the high signal-to-noise ratio provided by synchronous detection. This creates opportunities for applications in proximal sensing and dynamic phenotyping.

The framework also opens several directions for future research. Experimental validation of regulation fingerprints will be essential for testing the predictive power of the model and for constraining or falsifying alternative mechanistic descriptions of photosynthetic regulation. Extending the analysis to additional pathways, such as cyclic electron transport, state transitions, or metabolic feedbacks, may reveal further spectral signatures and richer multi-timescale dynamics. Combining frequency-domain measurements with spatially resolved sensing could likewise provide insight into how regulatory processes operate across tissues, canopies, and fluctuating natural environments.

More broadly, these results suggest that fluctuating light, often regarded as experimental noise or an environmental complication, can instead be used as a structured probe of photosynthetic function.

Frequency-domain analysis, therefore, provides a general strategy for linking dynamic measurements to underlying biological mechanisms and for characterizing regulatory processes through their spectral signatures. This perspective transforms fluctuating light from a source of variability into a quantitative tool for exploring the structure, dynamics, and regulation of photosynthesis.

## 5. Methods

### 5.1 BDM2 model framework

The simulations employed the BDM2 model introduced by Niu et al. (2025), which describes photosynthetic electron transport, energy balance, and nonphotochemical quenching using a minimal dynamical framework. The model consists of six ordinary differential equations representing the redox state of the plastoquinone pool, PQ(t), the redox state of the photosystem I donors, PI_ox_(t), the lumen proton concentration, H_L_(t), the ATP concentration, ATP(t), the zeaxanthin content, Zea(t), and the concentration of protonated PsbS, PsbS_act_(t), all evolving during periodic light forcing.

The model architecture relevant to the present study is summarized in Fig. 8. It comprises a constitutive photochemical core, represented primarily by the coupled PQ–PI_ox_ dynamics, together with two NPQ modules describing fast PsbS-dependent and slower Zea-dependent regulation. Both regulatory pathways are activated by lumen acidification and feedback on the effective antenna size of PSII (σII), thereby influencing photochemistry and chlorophyll fluorescence. The separation between constitutive photochemical processes and adaptive regulatory feedback provides the mechanistic basis for the distinct high- and low-frequency domains identified in this work and enables their interpretation in terms of underlying biochemical processes.

**Fig. 8.**
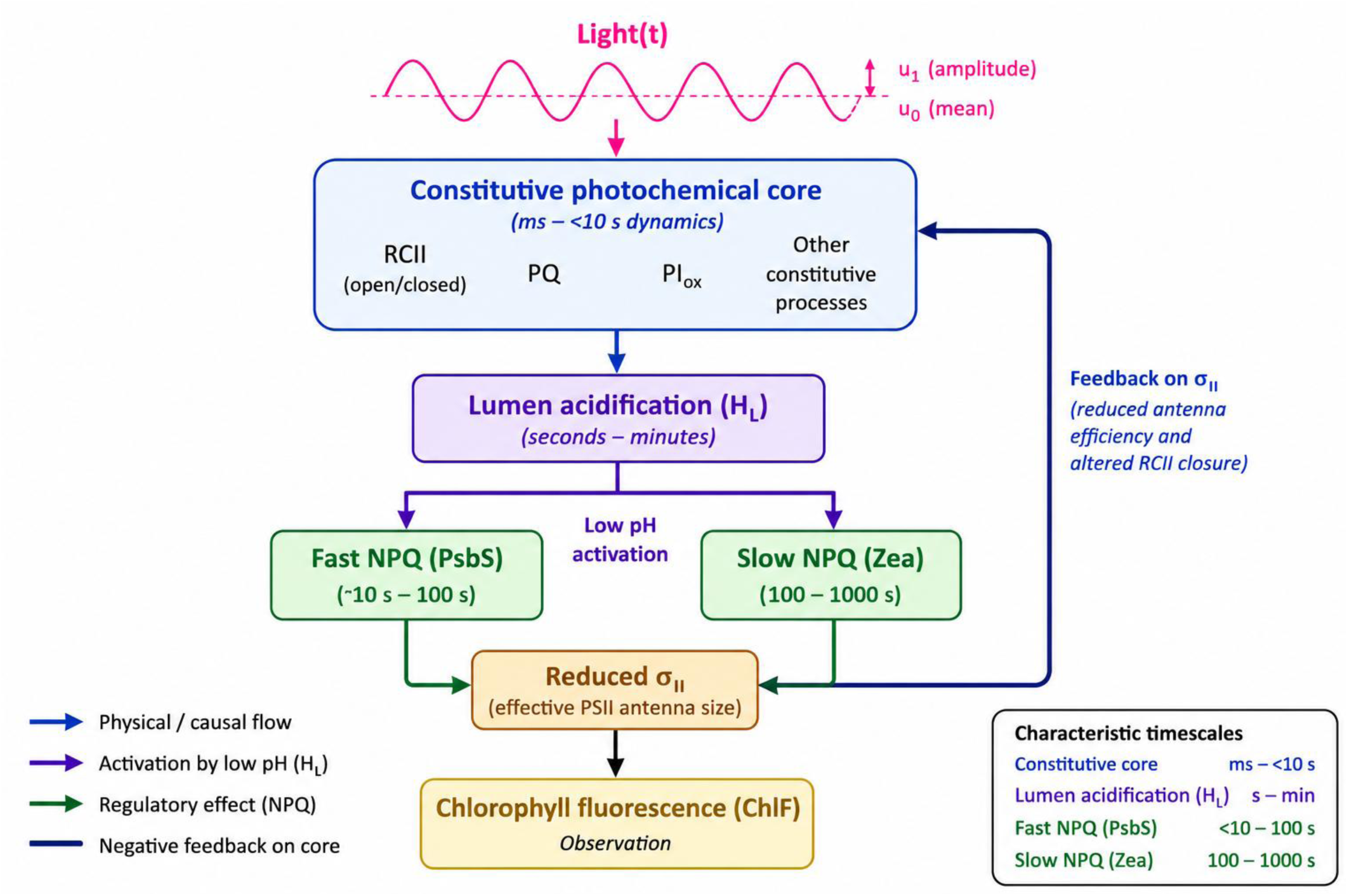
Simplified schematic representation of the BDM2 model. The model consists of a constitutive photochemical core coupled to fast (PsbS-dependent) and slow (Zea-dependent) nonphotochemical quenching (NPQ) pathways, both activated by lumen acidification (HL). The resulting feedback on the effective PSII antenna size (σII) modulates chlorophyll fluorescence (ChlF), providing the basis for the frequency-domain analysis presented in this work.

The activation and relaxation rate constants were 0.05 s⁻¹ and 0.004 s⁻¹ for PsbS and 0.01 s⁻¹ and 0.001 s⁻¹ for Zea, respectively. All other model parameters were adopted without modification from Niu et al. (2025).

### 5.2 Model-based interpretation of fluorescence dynamics

Following Niu et al. (2025), the normalized chlorophyll fluorescence signal is expressed as

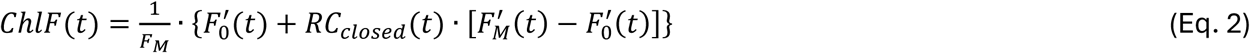

where 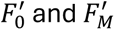 are the minimal and maximal fluorescence yield levels in the light-acclimated state, and RC_closed_ is the fraction of closed PSII reaction centers.

The quenched maximal fluorescence depends on the PsbS- and Zea-dependent components of nonphotochemical quenching,

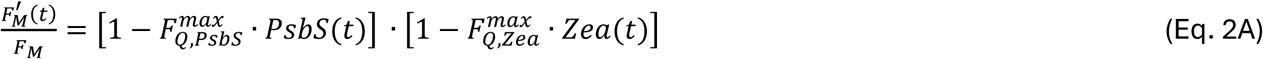

The corresponding minimal fluorescence level 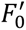 is related to 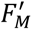 through the maximal photochemical efficiency Φ_max_ (Oxborough and Baker, 1997):

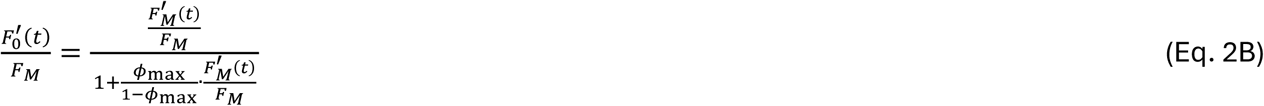

The fraction of closed reaction centers depends on light intensity, the redox state of the plastoquinone pool, and the effective antenna size of PSII,

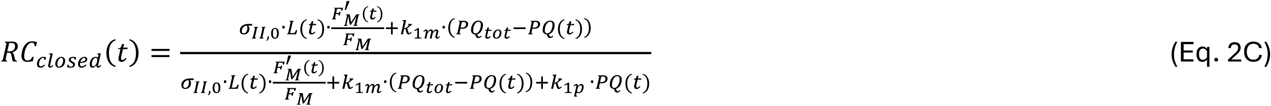

All parameter values were adopted from Niu et al. (2025).

Together, these expressions establish the mechanistic relationship between the measured fluorescence signal and the underlying photochemical and regulatory processes. In particular, chlorophyll fluorescence depends on both constitutive variables, such as incident light and the redox state of the plastoquinone pool, and on adaptive NPQ regulation mediated by PsbS and zeaxanthin. This separation provides the foundation for the linear decomposition developed in Section 5.3 and for interpreting frequency-domain features in terms of specific biochemical processes and feedback mechanisms.

### 5.3 Linear decomposition of the fluorescence response

To interpret the chlorophyll fluorescence responses mechanistically, the fluorescence signal was decomposed into contributions from the underlying processes. For small-amplitude oscillations within the linear regime, the fluorescence model (Eq. 2) can be linearized around the steady operating point, yielding

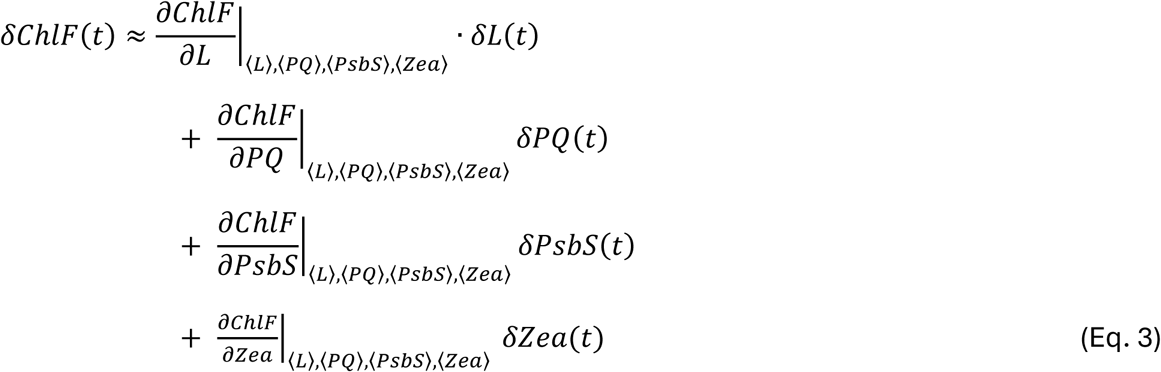

This expression separates the fluorescence response into four contributions: a direct light-driven term, a plastoquinone-mediated constitutive term, and two NPQ-related regulatory terms associated with PsbS and Zea. The decomposition provides the mechanistic basis for interpreting the frequency-dependent structure of the Bode plots and for linking specific spectral features to the underlying biochemical processes. In particular, it explains why the high-frequency domain is governed predominantly by constitutive photochemical dynamics, whereas the low-frequency domain reflects the activation of slower regulatory feedback mechanisms.

### 5.4 Transfer-function framework

In the linear time-invariant (LTI) regime, low-amplitude sinusoidal light modulation produces a response dominated by the fundamental harmonic at the forcing frequency. Under these conditions, the dynamic relationship between input and output can be described by a transfer function evaluated at the forcing frequency (Åström and Murray, 2008).

Let u(t) denote the light modulation and y(t) the response of an observable such as chlorophyll fluorescence. In the Laplace domain,

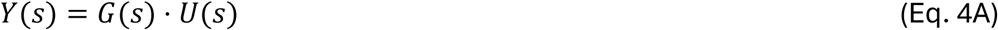

where *U*(*s*) and *Y*(*s*) are the Laplace transforms of the input and output signals, respectively, and *G*(*s*) is the transfer function.

For sinusoidal excitation within the linear regime, the input and output signals can be approximated as:

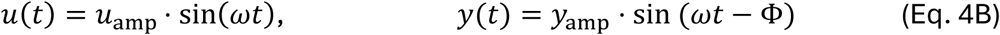

The corresponding frequency response along the imaginary axis is

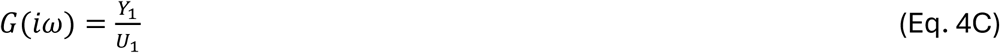

where *U*_1_ and *Y*_1_ are the complex Fourier coefficients of the fundamental harmonic of the input and output signals. The magnitude ∣ *G*(*iω*) ∣ defines the gain, and the argument arg*G*(*iω*) defines the phase shift between input and output.

By evaluating *G*(*iω*) over a range of frequencies, one obtains the frequency response of the system, commonly represented using Bode plots of gain and phase (see, e.g., Ingalls, 2013). In practice, *G*(*iω*) is estimated from the amplitude and phase of the fundamental harmonic of the measured response, as in Fig.2. Once higher-order harmonics become negligible (Fig. 4), the fundamental harmonic fully characterizes the local linear dynamics and provides a quantitative basis for frequency-domain system identification of photosynthetic processes.

### 5.5 High-frequency asymptotics of constitutive dynamics

In the high-frequency domain, chlorophyll fluorescence dynamics are determined primarily by the incident light and by variations in the redox state of the plastoquinone pool (PQ). Other state variables influence the response only indirectly through their effects on PQ dynamics or on slowly varying parameters entering the fluorescence model.

For the present parameterization, the constitutive dynamics of the BDM2 model are well approximated by an effective first-order lead–lag system,

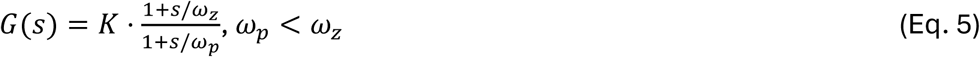

where *K* is the low-frequency gain and *ω_p_* and *ω_z_* denote the effective pole and zero frequencies, respectively. The gain approaches constant values in both frequency limits, whereas the phase shift vanishes at low and high frequencies and reaches a minimum at an intermediate frequency determined by the geometric mean 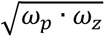.

The effective pole frequency is associated with the dominant relaxation mode of the coupled PQ–PI_ox_ subsystem around the light-acclimated operating point, whereas the zero frequency depends on the balance between the direct light-to-fluorescence feedthrough term and the indirect PQ-mediated response. Consequently, *ω_z_* is not a single kinetic rate constant but an emergent property of the coupled fluorescence–electron-transfer system.

At sufficiently high frequencies, the characteristic relaxation time of the PQ pool becomes much longer than the forcing period, so that *δPQ*(*t*)becomes negligible 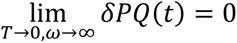 and the fluorescence response is dominated by the direct light-driven contribution. The corresponding asymptotic gain is therefore given by

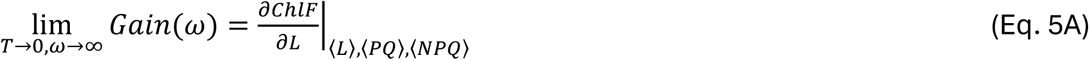

which, for the present model, yields

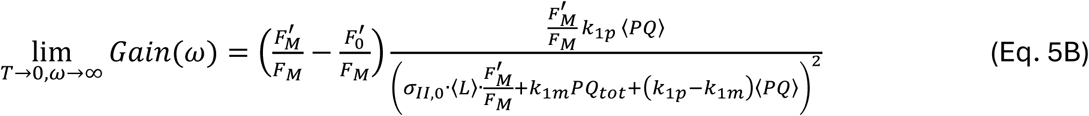

This asymptotic gain therefore reflects constitutive photochemical properties of the local operating point and is independent of slower regulatory feedback mechanisms. In principle, its value provides experimentally accessible information about the steady-state organization of the photosynthetic electron-transport system and may serve as a useful constraint for quantitative model identification.

### 5.6 Linearized PsbS dynamics and frequency-response features

The dynamics of PsbS activation in the BDM2 model follow Niu et al. (2025) and are described by

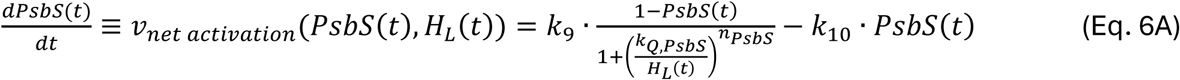

where *PsbS*(*t*) is the relative concentration of the active form of the PsbS quencher and *H_L_*(*t*) is the lumen proton concentration. The activation term follows a Hill-type dependence:

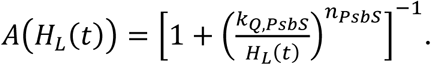

All parameter values were adopted from Niu et al. (2025).

For small-amplitude light oscillations, the system can be linearized around a steady operating point,

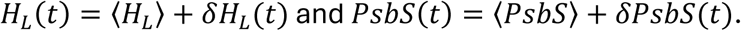

Linearization of Eq. 6A yields:

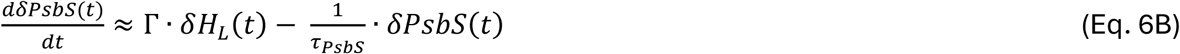

where

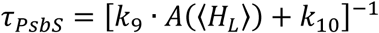

is the effective relaxation time and

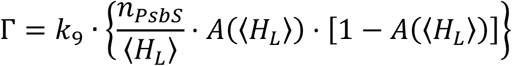

is the effective gain of the regulatory pathway.

The corresponding transfer function is

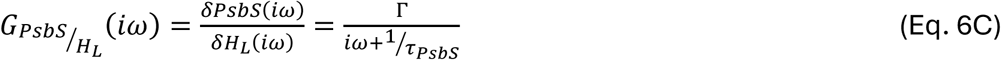

which is formally equivalent to a first-order low-pass filter.

The gain of this transfer function is:

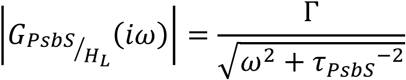

and the phase shift

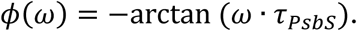

Thus, the relaxation time *τ_PsbS_*sets the characteristic frequency scale of the PsbS regulatory response and determines the frequency range over which gain transitions and phase extrema appear in the regulation fingerprints.

This explains the observed shift of phase maxima toward shorter periods with increasing light intensity (Fig. 7), since higher irradiance increases *A*(⟨*H_L_*⟩)and thus reduces *τ*_PsbS_.

The experimentally observable fluorescence signal is coupled to the regulatory dynamics through lumen proton concentration and PsbS activation. In the low-frequency domain (periods longer than approximately 10 s), constitutive processes are sufficiently fast that the lumen proton concentration follows the light modulation quasi-statically,

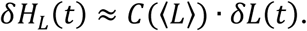

where *C*(〈*L*〉) depends on the steady-state operating point.

Under these conditions, the PsbS dynamics reduce to a first-order driven system with effective gain Γ and relaxation time *τ_PsbS_*. The gain parameter, therefore, determines the amplitude of the regulatory response and quantifies the sensitivity of NPQ activation to lumen acidification.

The transition between gain plateaus observed in Bode plots reflects the magnitude of Γ, providing, in principle, an experimentally accessible measure of the coupling strength between lumen proton dynamics and NPQ regulation.

The fluorescence signal is related to PsbS dynamics through

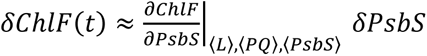 because the remaining contributions follow the slowly varying light oscillations without introducing substantial additional phase delays.

The dependence on PsbS arises through modulation of the maximal fluorescence level,

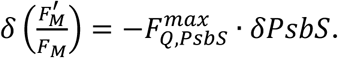

Thus, the phase shift observed in the ChlF signal directly reflects the delayed dynamics of PsbS activation, providing the mechanistic basis for interpreting low-frequency Bode-plot features in terms of NPQ kinetics. Within the linear decomposition of Eq. (3), the PsbS contribution therefore constitutes a distinct regulatory component that can be separated from constitutive photochemical dynamics.

Equivalent expressions apply to the slower Zea-dependent regulatory pathway.

## Acknowledgements

The BDM model and its code, used here for simulations, were generated by the HORIZON EUROPE EIC 2021 Pathfinder Open project DREAM, grant agreement no. 101046451and published in Niu et al. 2025. The author gratefully acknowledges Dušan Lazár and David Fuente for their critical reading of the manuscript and invaluable comments.

